# A deep learning model for analyzing noisy biological systems

**DOI:** 10.1101/2021.10.07.463577

**Authors:** Shangying Wang, Sara Capponi, Simone Bianco

**Author notes:** BAI Computational Innovation Hub, Altos Labs - Bay Area Institute of Science, Redwood City, CA.

## Abstract

Biological systems are inherently noisy so that two genetically identical cells in the exact same environment will sometimes behave in dramatically different ways. This imposes a big challenge in building traditional supervised machine learning models that can only predict determined phenotypic variables or categories per specific input condition. Furthermore, biological noise has been proven to play a crucial role in gene regulation mechanisms. The prediction of the average value of a given phenotype is not always sufficient to fully characterize a given biological system. In this study, we develop a deep learning algorithm that can predict the conditional probability distribution of a phenotype of interest with a small number of observations per input condition. The key innovation of this study is that the deep neural network automatically generates the probability distributions based on only few (10 or less) noisy measurements for each input condition, with no prior knowledge or assumption of the probability distributions. This is extremely useful for exploring unknown biological systems with limited measurements for each input condition, which is linked not only to a better quantitative understanding of biological systems, but also to the design of new ones, as it is in the case of synthetic biology and cellular engineering.

## 1 Introduction

Synthetic biology is a scientific discipline which aims at creating new biological parts and structures or modifying existing one, often with a specific application or objective. A major drawback of synthetic biology is the inherent difficulty of the process of designing genetically-encoded biological systems in a more predictable and programmable way. The relationship between the genotype, defined as the set of genetic information encoded in the DNA, and the phenotype, defined as the macroscopic realization of that information, is still unclear. The emergence of a specific phenotype may be linked not only to gene expression, but also to environmental perturbations and experimental conditions. Building a biology system is, in other words, like building a biological actuator, capable of yielding a specific phenotype by varying the genetic and environmental/experimental factors.

Phenotypic variation is ubiquitous in biology and is often traceable to underlying genetic and environmental variation. However, identical genotype and environmental conditions are not sufficient to guarantee a unique phenotype. This is mainly due to the inherent stochastic nature of biological processes. Random fluctuations may alter the levels of the biochemical components and drive the system to a specific phenotype. For example, in the case of protein synthesis, the bursty nature of messenger RNA (mRNA) expression causes the expression level of proteins to always fluctuate. Proteins typically have longer lifetimes than these random “bursts”, which results in a buffering effect, thus ensuring biochemical robustness of cellular homeostasis. Moreover, noise in the expression of a given gene can propagate to generate additional noise in the expression of downstream genes. Furthermore, many important cellular processes rely on actual physical interaction and hence spatial proximity between reactants (for example, RNA polymerase and DNA) and other physical criteria which, given the diffusive dynamic nature of the cell, occur stochastically. Noise is a key component of a well functioning biological system, and plays an essential role in key cellular activities 12, 4, 10, 15, 8. However, the nature and effect of noise in biological systems is still poorly studied and hence partially understood, which limits the use of machine learning tools in biology.

Particularly, building machine learning tools to elucidate the relationship between underlying genetic and environmental conditions to phenotypic observations is a challenge. One critical deficiency of this type of machine learning predictors is the general inability to call a given observation as an outlier relative to the data that the model has been trained and tested on 2. More specifically, there is a distribution of outputs (all possible phenotypic values) corresponding to each unique set of inputs (genetic and environmental condition).

A naive deep learning predictor will make a prediction based on the mean of all available observations for each unique set of inputs. This is problematic in the case of stochastic processes in the presence of limited observations, where the mean value cannot represent the entire dynamics of the system. Based on the central limit theorem in statistics, the sampling distribution of the mean for a variable will approximate a normal distribution given a sufficiently large sample size. The sampling distributions of the mean cluster more tightly around the population mean as the sample sizes increase. Conversely, the sampling distributions of the mean for smaller sample sizes are much broader. For small sample sizes, it is not unusual for sample means to be further away from the actual population mean. Furthermore, even with a very large sample size and a reliable machine learning predictor, the sample mean cannot provide insightful information for specific inputs because the whole probability distribution is important in understanding the whole characteristic of the population along with the average.

Figure 1 reports a simple examples of the difficulties in correctly mapping the inputs to the outputs if limited measurements are obtained. Let us assume that there are two different inputs (Input_1_ and Input_2_) to the biological system. For each input, experiments are performed and some measurements are taken. Figure 1 shows the noisy time course of a desired phenotype with 3 repeated measurements for each input. We also show the histogram of the steady-state value with 100 and 10000 measurements. The red line represents the sample mean. As one would expect, the larger the sample size, the smoother the histogram. Based on the histogram, it is possible to conclude that the output of Input_2_ is different from that of Input_1_, with the histogram in the former case shifting to the left. As well known, the analysis of small datasets may not allow to correctly capture the different phenotypes relative to Input_1_ and Input_2_. Moreover, in this toy example, we have assumed the probability distribution of the output to be Normal. However, for real biological systems, the type of probability distribution can be quite different depending on the inputs, which can further complicate the inference.

**Figure 1:**
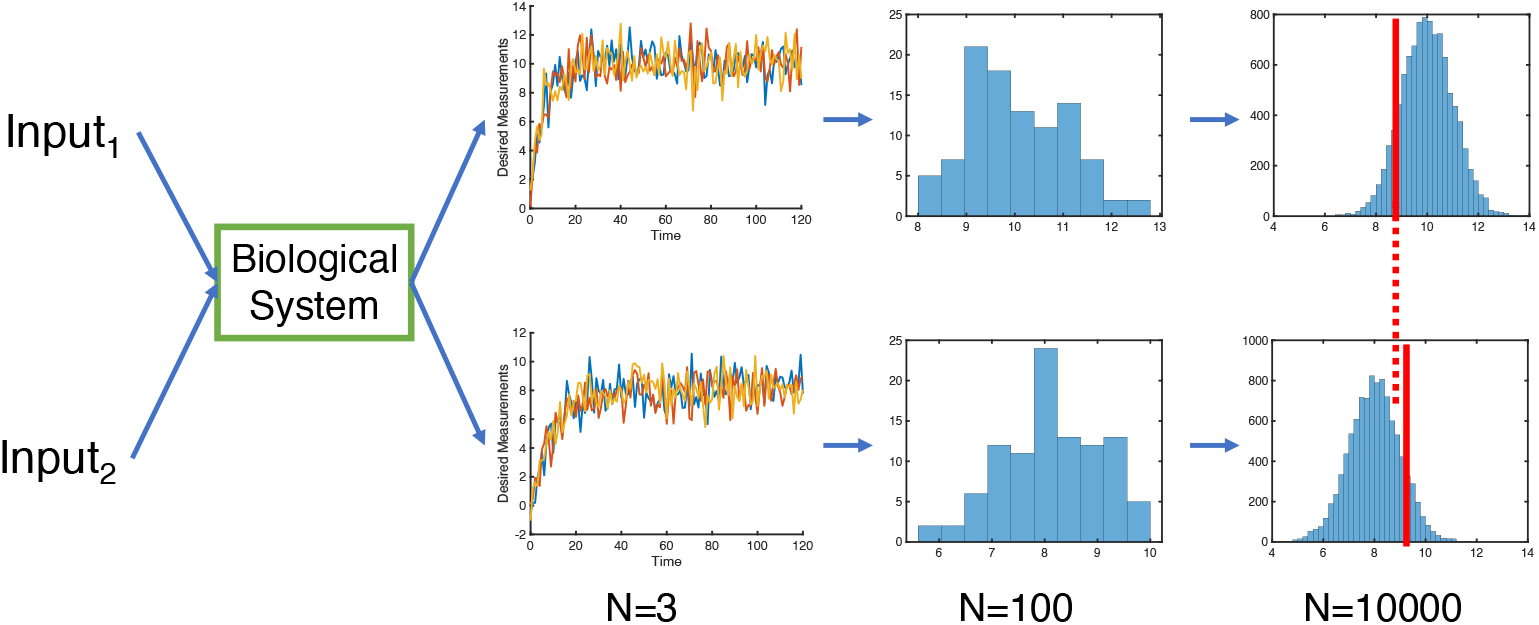
Mapping between inputs and outputs for a biological system. In this example we have used a Normal probability distribution for the output. The figure shows 3 time courses for two different inputs (Input_1_ and Input_2_) of the biological system respectively, and the histogram of the steady state value with 100 and 10000 measurements.

The aim of this work is to introduce a machine learning method capable of inferring the conditional probability distribution *CPD*(*y*|*x*) of the output variable *y*, conditional on inputs *x* without prior knowledge or assumption of the distribution itself.

## 2 Related work

### 2.1 Predictive machine learning algorithms to infer the sample mean with reduced noise

There has been little work devoted to using machine learning algorithms to increase the prediction accuracy of the mean value of a population in the presence of few noisy observations. Among these, we cite ensemble techniques, distance-based algorithm and single-learning-based techniques to identify noisy instances 6. However, knowing the whole probability distribution is still fundamentally needed to understand the characteristic of the biological phenomenon.

### 2.2 Inferring probability distribution vs noise reduction with increased sample size

Probability distributions for each input condition, i.e., conditional probability distributions, are generally estimated when sufficient observations are collected, through, for example, experiments or stochastic simulations 13, 9. Such methods are often expensive, time consuming or both.

If the conditional probability distribution of a target variable for a small subset of input combinations can be estimated, it has been shown that using a recurrent neural network, like a Long Short-Term Memory (LSTM) network, it is possible to predict the distribution of a larger set of inputs 17. In fact, the algorithm can be used to explore the whole input parametric space, as it has been shown for the myc-e2f pathway in cell-cycle progression 17, with high accuracy of prediction.

Absent a sufficiently large sample size, the estimation of the conditional probability may not be possible. With this work, we aim at solving this problem. Figure 2 is a schematic plot describing the inference of conditional probability distributions from limited observations. With varying input conditions, such as temperature, growth rate, lysis rate, etc., for each set of inputs, limited observations can be obtained.

**Figure 2:**
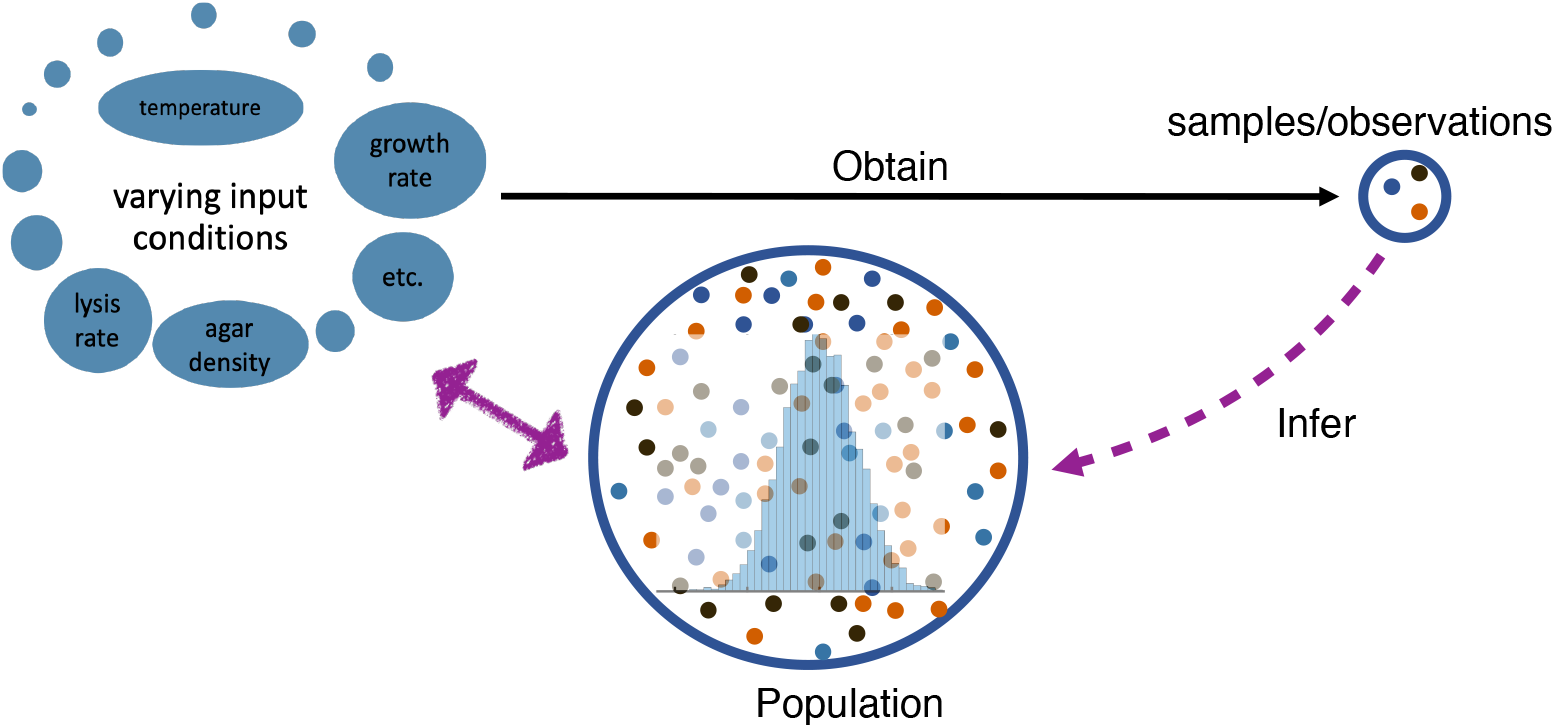
Research goal: infer the conditional probability distribution from few observations.

Moreover, it is often common to assume a probability distribution for the variable of interest 1 when it is unknown, through various ad hoc assumptions on the nature of the data. For example, for continuous and effectively unbounded outputs (within the region of probable values), a Gaussian distribution is typically used; For continuous and positive outputs, a Weibull distribution can be used. For discrete outputs (only integer values are allowed) and have a lower bound of zero, a negative binomial distribution might typically used.

Indicating the probability distribution assumed for the input data with the function *P* (*y*|*θ*), where *y* represents all possible values of the target variable and *θ* represents the hyperparameter(s) which set the exact shape of the distribution, simple methods exist to predict the value(s) of *θ*, which then allow the probability distribution function *P* (*y*|*θ*) to be calculated. In many cases, however, it is not possible to make educated or informed guesses about the shape of the distribution. In the next section, we introduce a machine learning framework capable of inferring the probability distribution of the output of a biologic system in the absence of prior assumptions about the shape of the distribution and using a limited set of observations per given biological input.

## 3 System and methodology

### 3.1 A simple gene regulatory network

Messenger RNA (mRNA) transcription is an essential process in protein synthesis. It is mediated by the presence of promoters and other elements which initiate and regulate the process. A conceptual model of the most elementary multi-state promoter, namely the two state promoter, is shown in Figure 3. The gene itself randomly transitions between transcriptionally inactive “off” state and active “on” state in “bursts” of transcriptional activity. There are four parameters: *k*_*on*_, the rate at which the gene transitions from the inactive to the active state, which determine the rate of transcription bursts; *k*_*off*_, the rate at which the gene transitions from the active to inactive state, which determines the duration of transcription bursts; *v*, the rate of transcription when the gene is in the active state, which determine the burst size, i.e., how many mRNAs are produced during each burst; and *δ*, the rate of mRNA degradation. This process was first modelled and analyzed by Peccoud and Ycart 11. We choose to apply our framework on this system because it is simple and has an analytic solution of the probability density function of mRNA (Eqn. 1), and therefore allows for direct validation of our method 16. The conditional probability density can be written as follows:

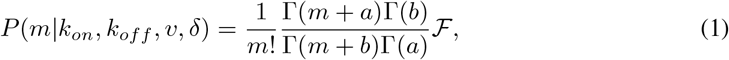

where 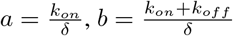 and 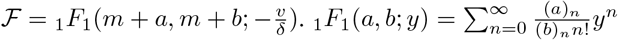 is the confluent hypergeometric function of the first kind, where (*a*)_0_ ≡ 1 and (*a*)_*m*_ ≡ *a*(*a* + 1)(*a* + 2)…(*a* + *m* − 1).

**Figure 3:**
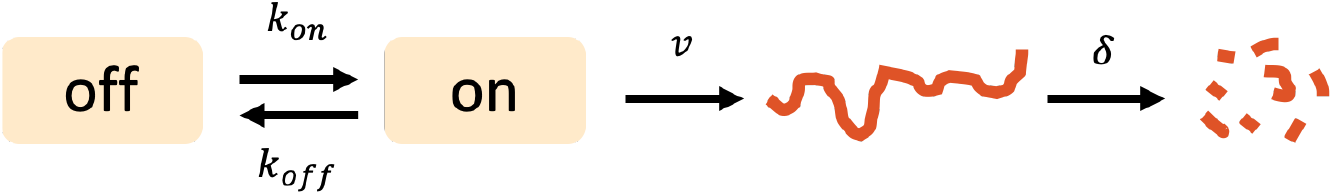
Scheme of a simple gene regulatory network. The gene randomly transition from an *on* to an *off* state and viceversa, with rates *k*_*on*_ and *k*_*off*_, respectively. The gene can transcribed in the *on* state at a rate *v*, and degrades at rate *δ*.

Equation (Eqn. 1) has only three dimensionless parameters which directly affect the outcome: 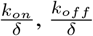 and 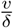. In what follows, we fixed *δ* = 1 and only vary input parameters *k*_*on*_, *k*_*off*_ and *v* for simplicity. Experimental observations have indicated that the half-life of the mRNA is on the order of 5 − 10 minutes in bacteria 14, around 10 hours in human 18. Any scaling will result in the same stationary distribution, as such distributions are, by definition, independent of time.

For this model, there is not only intrinsic noise, but also measurement noise. The measurement noise is due to the restriction that all the measured mRNA numbers shall be integers.

Figure 4 illustrates that even for this simple gene regulatory system, with different parameter values, the probability distribution can take various forms. It can sometimes be monotonically or exponentially decreasing, bimodal, a step function or a Gaussian. Thus we cannot use a pre-assumed distribution type approach for this system.

**Figure 4:**
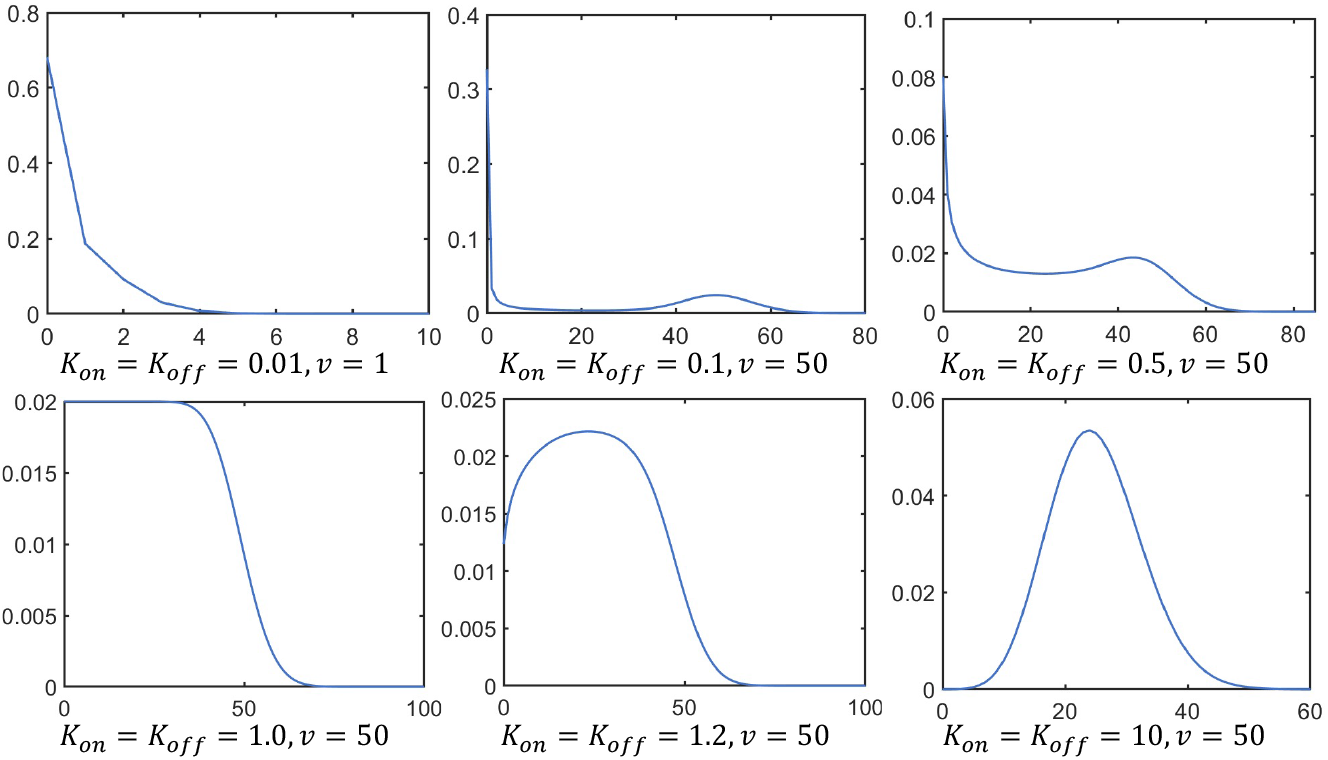
Examples of theoretical probability distributions for the biological system of Figure 3.

### 3.2 Data preparation

For the system described above, we randomly generate 13000 parameter combinations by tuning *k*_*on*_, *k*_*off*_ and *v*. (0.01 *< k*_*on*_, *k*_*off*_ *<* 1000 and 0.01 *< v <* 200) 10. 10000 parameter sets are used for the training and 3000 are used for testing. We calculate the probability distribution for mRNA number from 0 to 300 for each parameter combination to estimate neural network performance after training.

We then numerically computed the cumulative distribution function (cdf) for the system based on Eqn. 1. For any random variable, the cdf is always a non-decreasing function with 0 ≤ *cdf* ≤ 1. Thus, by drawing a random number *r* (0 ≤ *r* ≤ 1) and then by finding the corresponding inverse value on the abscissa, a sample value of random variable can be generated. This method is called inverse transform sampling 3.

By using the inverse transform sampling method, we randomly draw *n* samples (*n* = 1, 3, 10 and 100) for each parameter combination to represent the acquisition of a subset of the observations, which are used in the training process to minimize the loss function.

### 3.3 Machine learning framework

Classic regression models would take a set of inputs *X*, and train machine learning models to predict the value of the target variable *Y*. The value is usually a single observation value, or a mean value based on a few observations. The loss function is chosen to minimize the neural network predictions with the mean value of the observations. In this study, we present a Recurrent Neural Network (RNN) based model for the data-driven inference of the conditional probability function of noisy biological systems. As mentioned, our RNN model aims at directly predicting the probability distribution function without any prior knowledge or assumption, except for continuity.

In this work, we used LSTM Neural Networks implemented in Python 3.7.8 and TensorFlow v2.4.1 without GPU implementation. LSTM, which is a special kind of RNN, is proposed to overcome the difficulties in learning a long-term dependency structure 7, 5. LSTM uses multiple gating functions to conserve the information stored in its internal state for a longer period of time. The neural network architecture is illustrated in Fig. 5.

**Figure 5:**
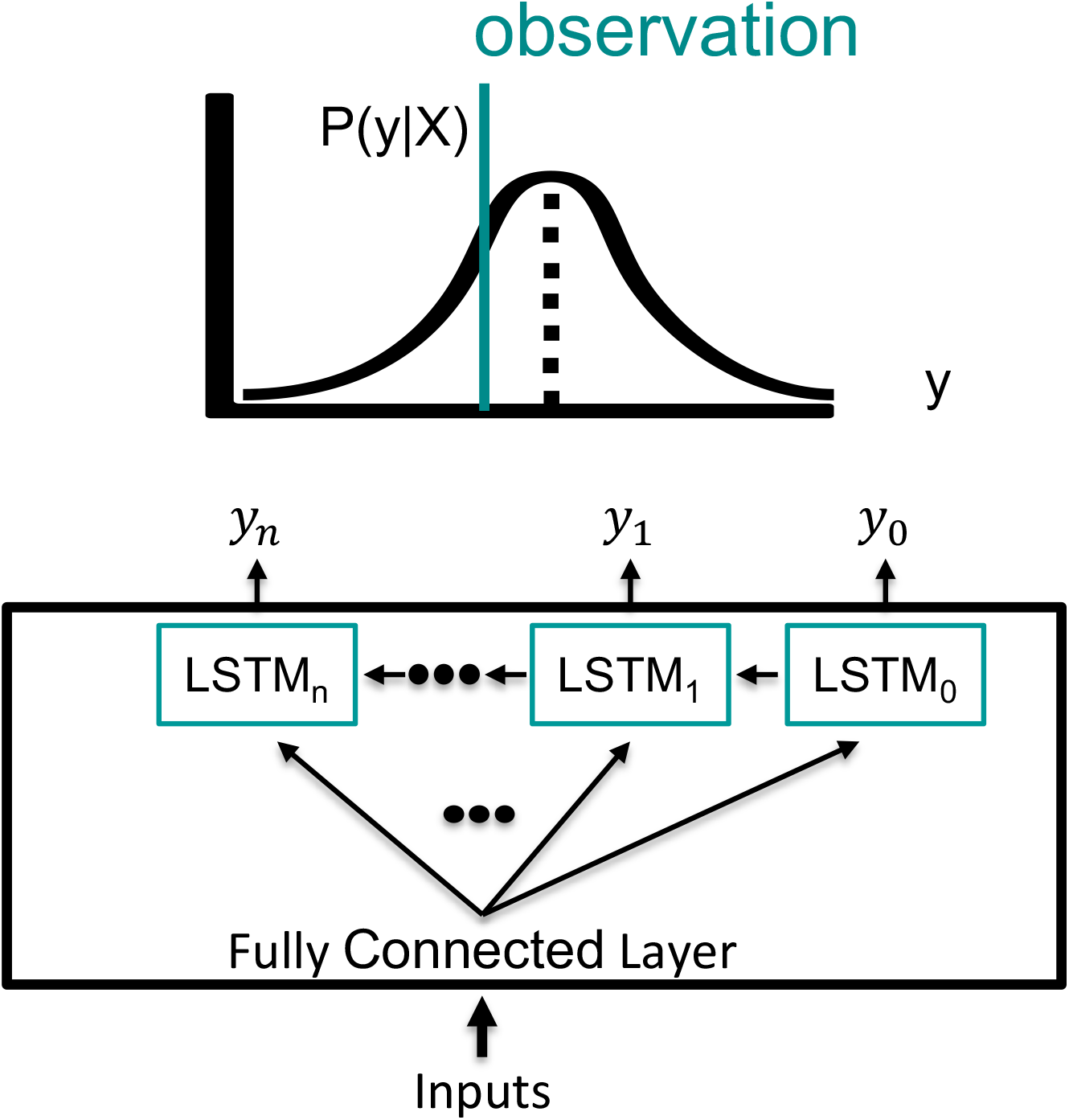
Machine learning framework.

Our model also employs a negative log-likelihood function as the loss function (Eqn. 2). The loss function is trained with limited output observations (e.g., the number of mRNA produced). The framework can be applied to any biological system (e.g., a cell) whose input parameter space is large and virtually impossible to explore experimentally (e.g., genetic modifications, physical modifications, environmental modifications), and whose output is a desired phenotype (e.g., cellular growth, cellular adhesion, cellular sensing).

The use of a RNN with a negative likelihood loss function has never been attempted to estimate an unknown probability distribution of a biological phenotype. The combination of the minimization of the loss function using observations (which are not a ground truth), and the training of the RNN using input parameters is also non-obvious.

Suppose there is a data set *D* = {(*y*_*i*_, *x*_*i*_); *i* = 1, …*M*}. The parameters of the artificial neural network Θ are estimated by minimizing a negative log likelihood function *l*(Θ).

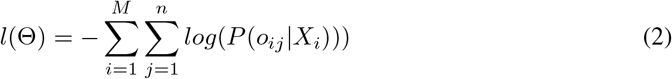

where *M* is the data size, *n* is the number of observations, *X*_*i*_ represents the *i*th input parameter combination, *o*_*ij*_ represents the *j*th output observation corresponding to the *i*th input parameter combination, and *P* is the probability function of the observations to the parameters, which is first initialized to be uniform during the neural network initialization process. The loss function is trained using a limited set of collected observations in order to remove the uncertainties that are inherent in a RNN. Its purpose is to maximize the probability that the observed data is within the predicted probability distribution of the RNN. It does this by calculating the sum of all of the probabilities of the observed data within each parameter. Because the loss function is a negative log-likelihood, its integration into the RNN minimizes the loss of observations that would otherwise fall outside of a typical mean analysis. The first step in the training of the RNN described herein is the establishment of the loss function. During training of the RNN, the loss function compares the prediction outcomes to the desired output resulting in output values throughout the time series and propagation of the loss function back through the RNN to update the input weights; thus, every node that has participated in the calculation of the output associated with the loss function has its weight updated to minimize the error throughout the RNN.

### 3.4 Pipeline

#### 3.4.1 Data collection/generation/cleaning

We assume that data is collected either through experiments or *in silico* for various input conditions if a stochastic formulation the biological system is available. Input values can be chosen randomly. Our methods accepts normalized to a standard scale input values. If the observations are discrete numbers (mRNA counts, protein counts, etc.), they need to be integer values, and are included as one of the predicting points for the neural network (NN) outputs. If the observations are continuous values (concentrations, optical density measurements, etc.), they formally need to be bound from above to ensure that observations are within the prediction range of the neural network.

#### 3.4.2 Neural network construction

The neural network input layer inputs all the input conditions. The neural network output layer outputs the probability distribution. If the observations are discrete numbers, the neural network predict the probability value for all possible discrete numbers and a Softmax layer is implemented to ensure that the sum for the predicted probability for any input condition equals 1. If the observations are continuous values, it is necessary to discretize the possible observation range into bins. The neural network is only predicting the probability value for the center of the bins. The probabilities for other values are interpolated. The last layer is normalized with a normalization factor to make sure the cumulative trapezoidal numerical integration of the probability distribution equals 1.

#### 3.4.3 Neural network training

Neural network nodes are randomly initialized. The output probability distribution are initialized to a uniform distribution and modified each training epoch to minimize the loss function. The gradient was clipped to prevent exploration. The neural network is trained using a training data set consisting of the input conditions and one output observation. To test the performance, measurements of a small batch of data is repeated multiple times to estimate a sample probability distribution.

#### 3.4.4 Neural network predictions

After training, the neural network is ready to be used for predicting the probability distribution for any input. Trained neural network can be used to facilitate the quantitative understanding of biological systems as well as the design of synthetic gene circuits.

## 4 Results

### 4.1 Conditional probability distributions can be inferred with limited observations

The input of our algorithm is parameter combinations, specifically, *k*_*on*_, *k*_*off*_, and *v* for the proposed system. The output is the probability distribution of the intrinsically noisy observations. Figure 6 and Figure S.1, S.2, S.3 shows the neural network performance for the test dataset for sample size *n* = 1, 3, 10 and 100 respectively. The examples are randomly chosen from the test datasets. Table 1 listed the values of the loss function, *R*^2^ and the root mean squared error (RMSE) for the training and test dataset for *n* = 100, 10, 3 and 1 respectively. The accuracy is calculated by comparing the predicted probability distribution vs the real probability distribution. This is done after the neural network training is finished. Since the probability distribution is discretized into 301 points, the RMSE is calculated based on comparing (301 * *datasize*) values between the prediction and the ground truth. All the values are the average value from 3 neural networks.

**Figure 6:**
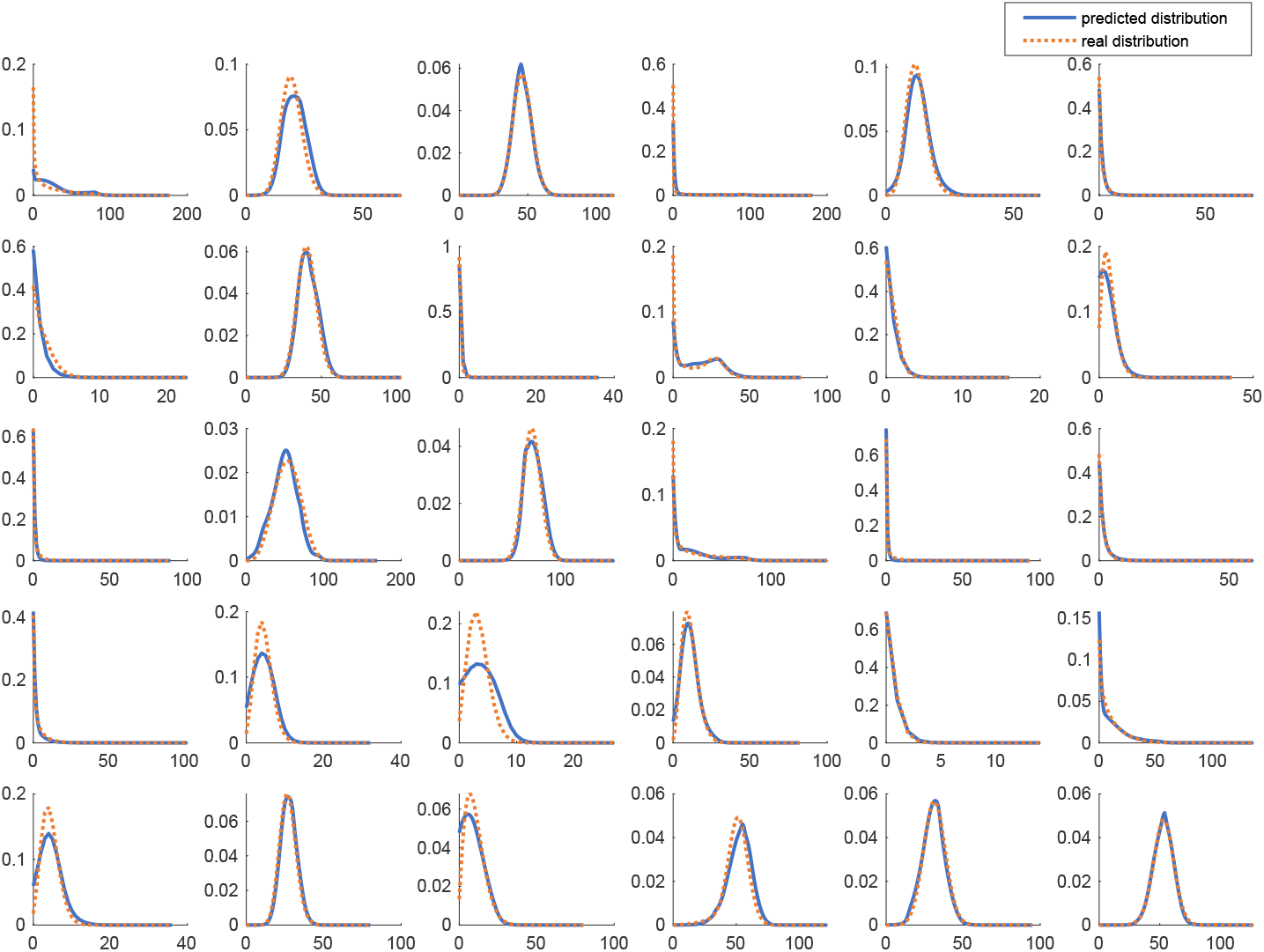
Neural Network performance of the test data for sample size n=1.

**Table 1:** Model Performance Comparison

| | $n = 100$ | | $n = 10$ | | $n = 3$ | | $n = 1$ | |
| --- | --- | --- | --- | --- | --- | --- | --- | --- |
|  | Train | Test | Train | Test | Train | Test | Train | Test |
| Loss | 1.40908 | 1.41247 | 1.40535 | 1.41159 | 1.40645 | 1.41636 | 1.40508 | 1.42919 |
| $R^2$ | 0.9768 | 0.9759 | 0.9762 | 0.9749 | 0.9770 | 0.9760 | 0.9735 | 0.9725 |
| RMSE | 0.00630 | 0.00641 | 0.00631 | 0.00643 | 0.00615 | 0.00628 | 0.00651 | 0.00662 |

The results in figure 6 and table 1 show that the model accurately predicts the probability distribution in various conditions and without any assumption, even when 3 or less observations per input condition are collected.

Using this method, repeated measurements under a fixed condition are not necessary to obtain a reliable estimation of the sample probability distribution, and resources may be devoted to explore the dynamic of the system in a wider range of parameter sets, which would allow to more easily detect different dynamical regimes, as well as give a reliable estimate of the variability of the outputs.

A consequence of the accuracy of the predictions of the distribution using deep learning is a higher accuracy in the estimation of the sample parameters and distribution momenta. As an example, in Figure 7 we report the sample mean and the mean predicted using the NN vs the real mean, calculated based on Eqn. 1). Larger sample size can largely reduce the noise and make the sample mean closer to the real population mean (top panel). By using our framework, the predicted mean calculated based on the predicted conditional probability distribution from the neural network can be estimated for significantly smaller sample sizes.

**Figure 7:**
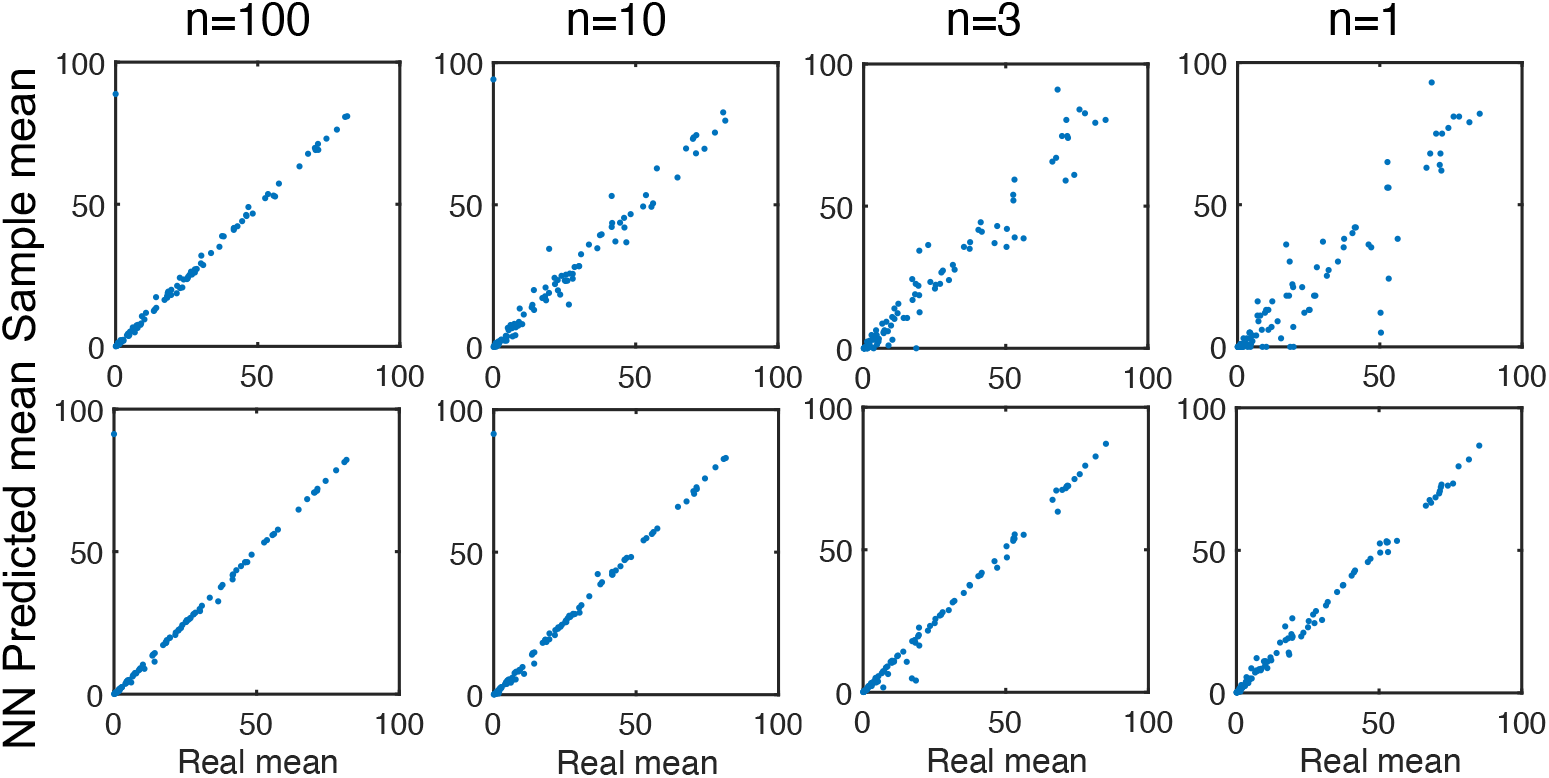
Sample mean (above), predicted mean (below) vs real mean.

## 5 Conclusion and future work

Biological processes are intrinsically stochastic, which often causes difficulties in our ability to investigate and understand biological data. Specifically, it hinders our understanding of the relationship between the underlying genetic, environmental conditions and phenotypic observations, which has important consequences for fields like synthetic biology and cellular engineering, where high reliability of the output prediction may be critical.

We develop a deep learning algorithm which is able to predict the probability distribution of the phenotype observations with as few as one observation per input combination. The key innovation of our study is that deep neural networks can be used to generate probability distributions based on few observation for each input condition, without prior knowledge or assumption on the form of the probability distributions.

Our approach overcomes the bottleneck faced when many observations per input condition are necessary to eliminate the effect of noise of phenotypic observations or reconstruct the conditional probability distribution of a stochastic process, which is often expensive and time consuming. It also reduces the barrier in implementing predictive machine learning model for “noisy” biological data.

## Code availability

The source code is available at github: https://github.com/syntheticbio/RNN_NLL.

## Acknowledgments

This material is based upon work supported by the National Science Foundation under Grant No. DBI-1548297.

## A Appendix

Due to the page limit, we include the neural network performance of the test data for sample size *n* = 3, *n* = 10 and *n* = 100 in the appendix.

**Figure S.1:**
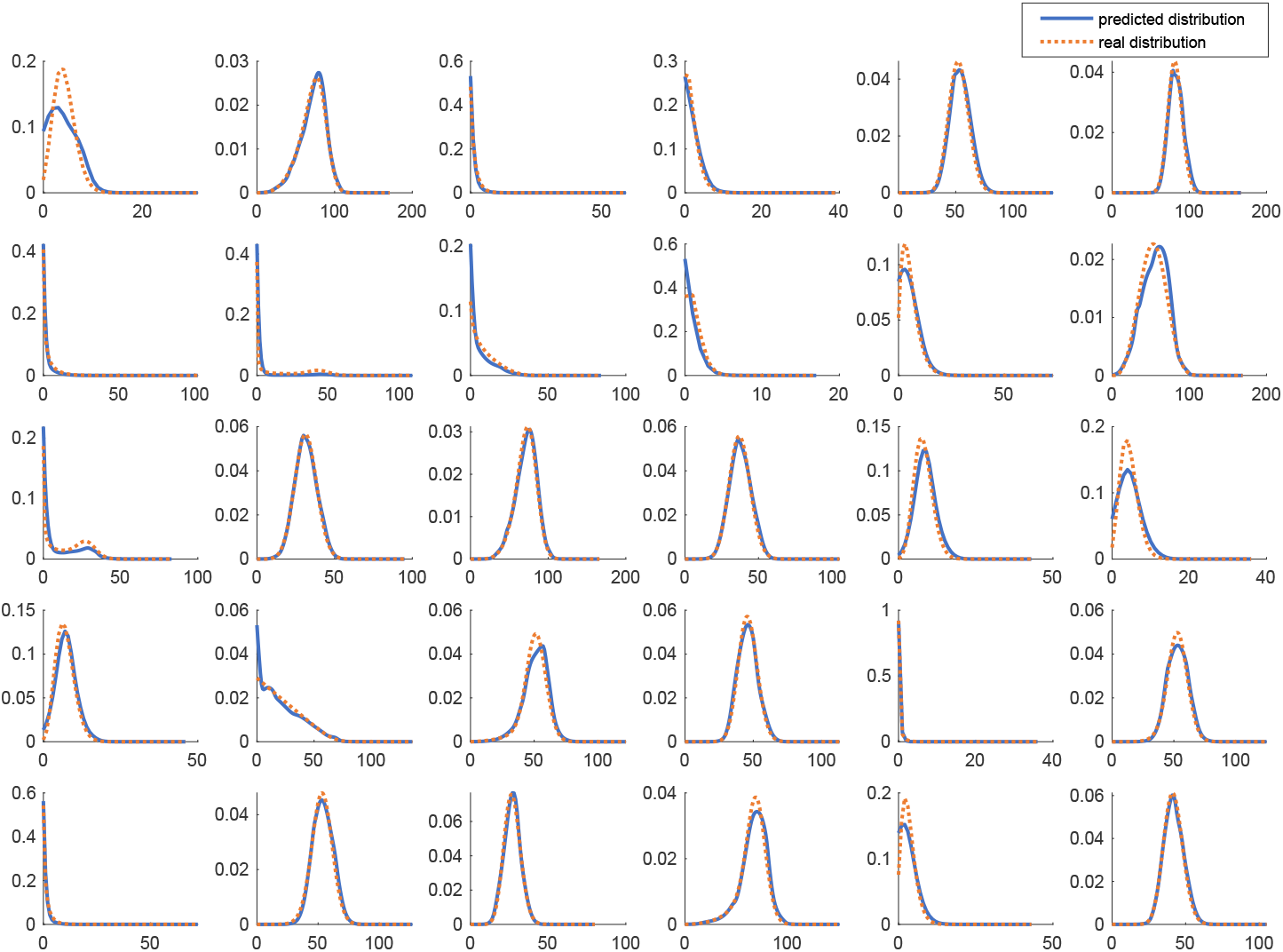
Neural Network performance of the test data for sample size *n* = 3.

**Figure S.2:**
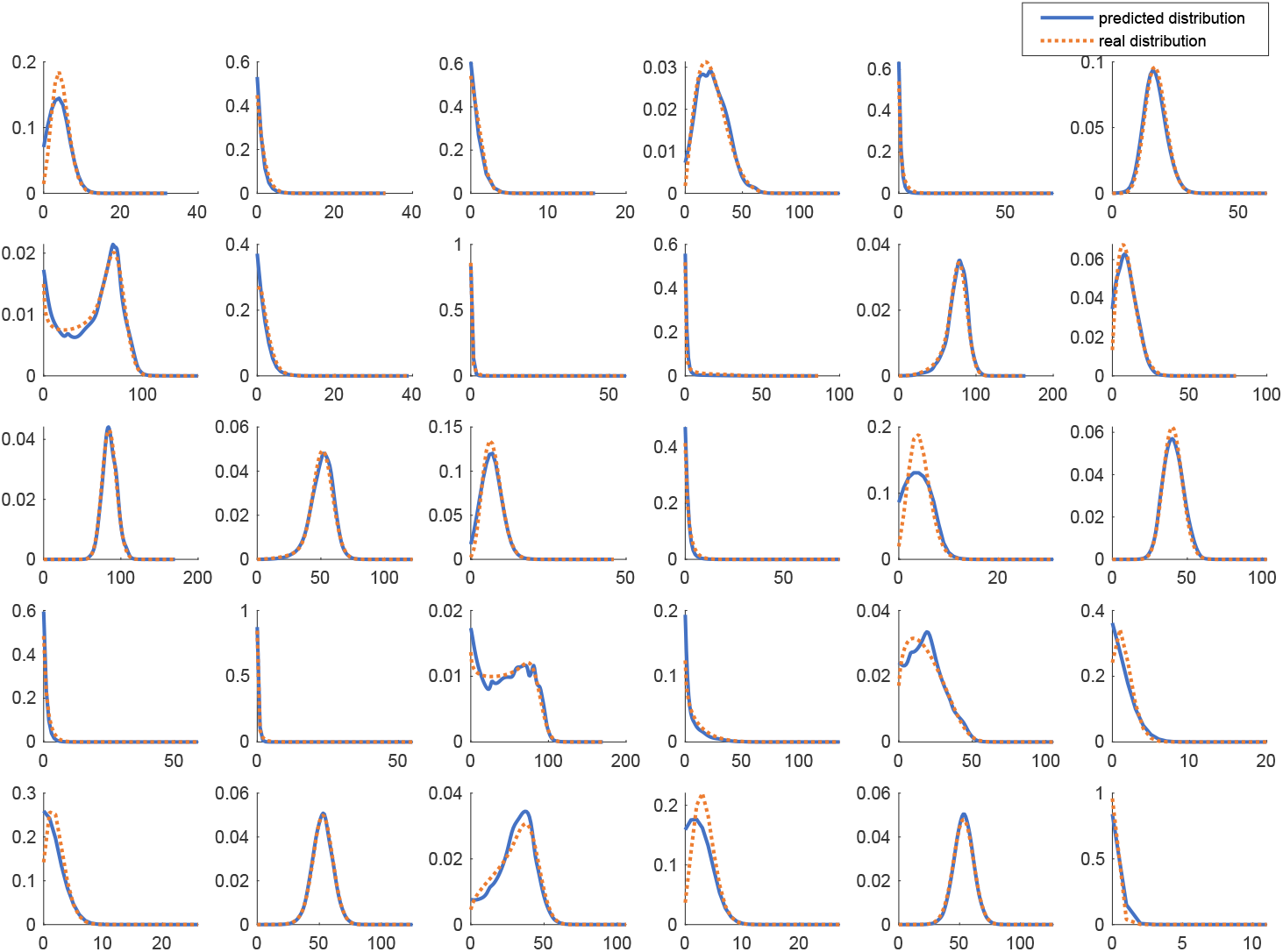
Neural Network performance of the test data for sample size *n* = 10.

**Figure S.3:**
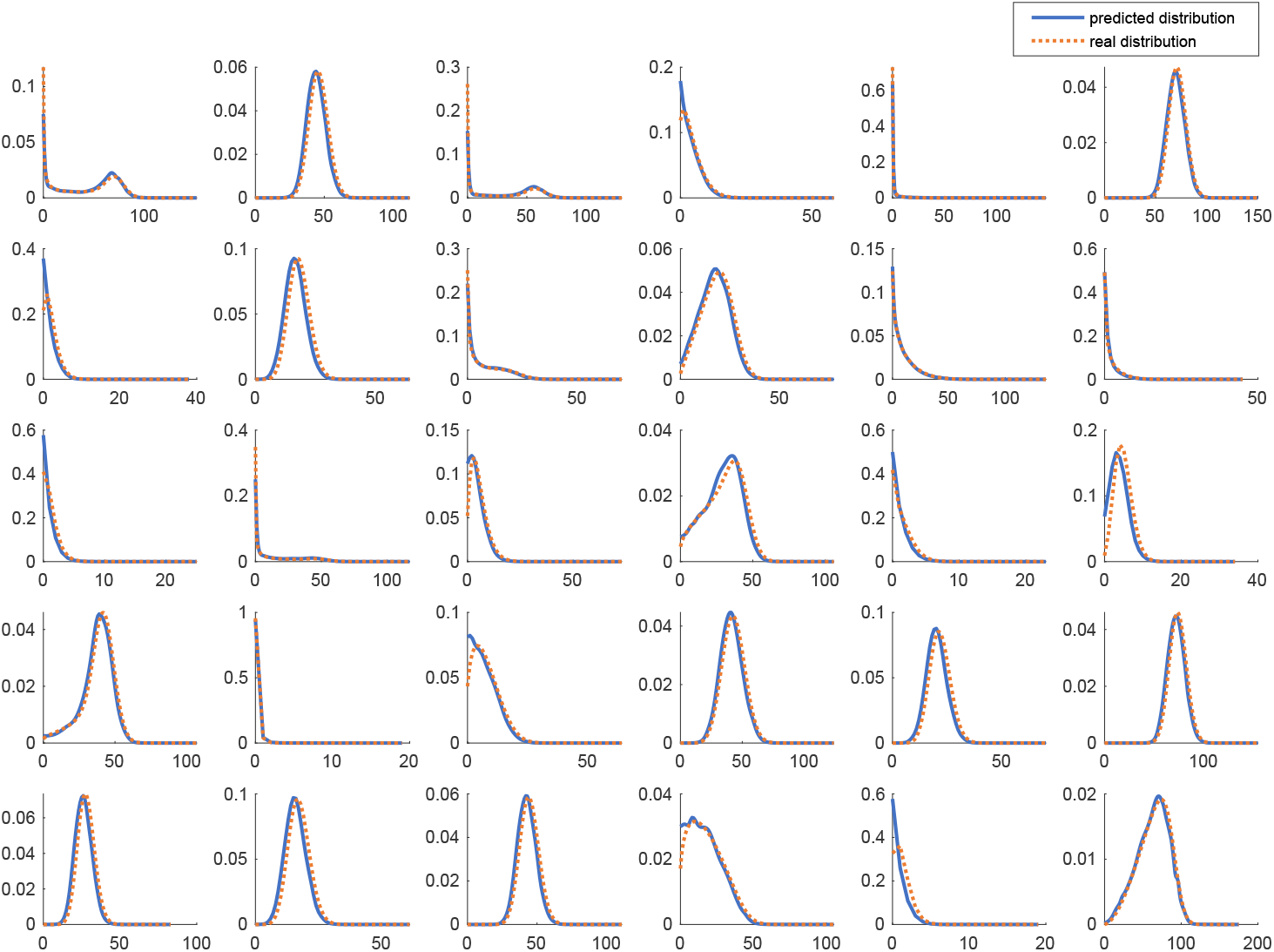
Neural Network performance of the test data for sample size *n* = 100.

